# Excessive sample reagent incubation time increases the risk of false rifampin resistance results in Xpert^®^ MTB/RIF Ultra tests performed on paucibacillary Mycobacterium tuberculosis sputum

**DOI:** 10.64898/2026.10.10.758171

**Authors:** Gayatri Shankar Chilambi, Kunj Modi, Paulami Rudra, Yiwang Chen, Soumitesh Chakravorty, David Alland

## Abstract

A high rate of false rifampicin resistance (RIF-R) was recently reported in paucibacillary sputum samples tested with the Xpert^®^ MTB/RIF Ultra (Ultra) assay. However, 96% of the studied facilities started the test 8 – 12 hours after addition of sample reagent (SR), exceeding the manufacturer’s recommended limit of 4 hours. Assessed whether excessive long SR incubation could lead to similar false RIF-R results in paucibacillary samples in a reference laboratory setting. We evaluated the performance of the Ultra assay following incubation of TB-negative sputum spiked with 300 CFU/mL and 100,000 CFU/mL levels of Mycobacterium tuberculosis (Mtb) H37Rv in SR for different durations (15 min to 24hrs.) at different temperatures (-20 to 37°C) to determine the effect on RIF result calls. No false RIF-R results were observed in these samples with 10^5^ CFU/mL Mtb at any of the conditions tested, even when they deviated largely from recommended limits. Similarly, freezing the 300 CFU/mL SR treated samples at minus 20°C for 18 h also prevented the development of false RIF-R results. However, sputum spiked with 300 CFU/ml Mtb, produced false RIF-R calls at longer incubation times and higher incubation temperatures than are recommended by the manufacturer. Approximately half of the results displayed melt patterns consistent with hetero-resistance, indicating SR-induced mutagenesis in a wild-type background. This study demonstrates that extended off-label SR incubation of paucibacillary specimens beyond the recommended 4-hour limit can generate false RIF-R results. Freezing paucibacillary specimens in SR may mitigate this risk although it is not yet an approved procedure.

## Introduction

Xpert^®^ MTB/RIF Ultra (Ultra) is a WHO-recommended low complexity automated nucleic acid amplification test (LC-aNAAT) approved for *Mycobacterium Tuberculosis* (MTB) and Rifampicin (RIF) resistance detection from sputum (1). Ultra was developed to overcome the shortcomings of its predecessor, Xpert® MTB/RIF (generation 4 assay or G4), by improving sensitivity to detect smear-negative and low bacillary load samples. A melting temperature-based analysis was introduced in Ultra to detect the *rpoB* Rifampicin resistance determining region (RRDR) mutations as opposed to G4 which detects the mutations based on a cycle threshold shift (delta Ct) between two probes. This delta Ct approach in some rare cases with paucibacillary specimens may lead to false positive RIF-R results due to Ct delays of probes D or E resulting from low bacillary load, leading to artifactually high delta Ct values(2). Other reasons for false positive RIF-R detection in G4 include the presence of silent *rpoB* mutations, or cross-reactivity with other NTMs (3).

According to WHO guidelines, the treatment procedure for a patient with false RIF resistant test result by a NAAT assay, but a RIF susceptible phenotypic DST (pDST) should be based on the NAAT assay results until additional tests such as sequencing, pDST on solid media or confirmatory testing on an alternative NAAT assay are performed (1). Consequences of detecting such false-positive rifampin resistance include incorrect patient treatment, unnecessary treatment with second line anti-TB drugs (3), side effects including hepatotoxicity, neurotoxicity (4) and increased healthcare costs (3). The melt-based resistance detection approach in Ultra assay was introduced to specifically mitigate false RIF-R results in the G4 assay, which were associated with (i) paucibacillary samples, (ii) silent mutations in *rpoB* RRDR or (iii) primer/probe cross-reactivity in samples with multiple NTM infections (2, 3)

Several recent studies have reported false RIF-R calls using Xpert^®^ Ultra, particularly in paucibacillary specimens, comparable to the trend observed in the G4 assay (2,8). One of these studies involved nationwide testing in Rwanda, where among 129 patients diagnosed as RIF-R TB cases by Ultra initially, only 32 % were confirmed as RIF-R, and the remaining 68% showed discordance on repeat Ultra testing, with 54 generating RIF-S results (5).These discordant Ultra results were specifically obtained with very low or low bacillary loads. It was reported that in 96% of the study sites, samples were loaded into the Ultra carts between 8 -12 h post-addition of SR, although no specific information was available on the incubation times of individual SR/sample mixtures for the 54 false RIF-R cases (5).

Previous studies which evaluated SR incubation of sputum beyond the recommended limit of 15 minutes to 4 hours (3) have shown that prolonged incubation in SR leads to deamination of DNA due to the presence of sodium hydroxide (6, 7). These studies were performed with the G4 assay. However, no systematic study has been performed to evaluate the effect of prolonged SR incubation on the Ultra assay and its consequence on RIF-R calls. Here, we perform a systematic analysis of prolonged SR incubation of contrived sputum samples at different time and temperature conditions which significantly deviated from the manufacturer recommended conditions for incubation in SR. We evaluated the SR incubation protocol deviations using contrived sputum samples representing both high and low bacillary loads to assess whether this may be related to the excessive false RIF-R calls observed in the Rwanda study.

## Materials and methods

### Sample preparation and testing procedure

To prepare contrived sputum samples for testing, we used quantified stocks of an attenuated version of the *M. tuberculosis* H37Rv mc^2^6230 laboratory strain prepared and quantified by CFU plating as described previously (2, 8, 9). Multiple aliquots of the same quantified stock were used to perform the entire experimental procedure to avoid any operator induced CFU variations, which may happen while using different cell stock preparations.

We evaluated the performance of the Ultra assay following prolonged incubation of TB-negative sputum spiked with lower (300 CFU/mL) and higher (100,000 CFU/mL) loads of *Mycobacterium tuberculosis* H37Rv in Xpert^®^ sample reagent (SR). The spiked sputum samples were incubated in SR for different durations and at different temperatures to determine the effect on RIF-R results. We assessed the Ultra results for samples subjected to SR-incubation at 37°C (at hold times of 15 minutes, 1 h, 2 h, 3.5-4 h, 9 h, 16 h and 23.5-24 h); 30°C (at a single hold time of 3.5-4 h only); 25°C (at hold times of 3.5-4 h, 9 h, 16 h and 23.5-24 h); 4°C (at hold times of 9 h, 16 h and 23.5-24 h) and −20°C (at a single hold time of 18 h only) for the contrived sputum samples containing 300 CFU/ml Mtb (Fig 1). The samples spiked with 100,000 CFU/mL Mtb. were incubated in SR at three different temperatures of 37°C (at hold times of 3.5-4 h and 9 h), and at 4°C and 25°C with a single hold time of 9 h at both temperatures. The sputum-SR mix was added to the cartridge sample chamber only after the incubation for the specified duration was complete. Xpert Ultra carts were loaded into the GeneXpert instrument after the SR incubation, and tests were run following the manufacturer’s instructions. A minimum of 30 replicates were run for each condition tested, amounting to a total of 120 replicates with the samples containing 10^5^ CFU/ml Mtb, and a total of 480 replicates with the samples containing 300 CFU/ml of Mtb (Table 1), since more conditions were tested for samples with a lower concentration of Mtb. Mean IS*6110*/IS*1081* and earliest *rpoB* Ct values, melting temperatures (Tm), and result outputs were assessed for each condition. The first derivative fluorescence data used to plot the melt curves were extracted from the. nxx run files. Percentage of false RIF-R was calculated by dividing the total number of RIF-R calls by the total number of valid RIF-S calls.

**FIG 1.**
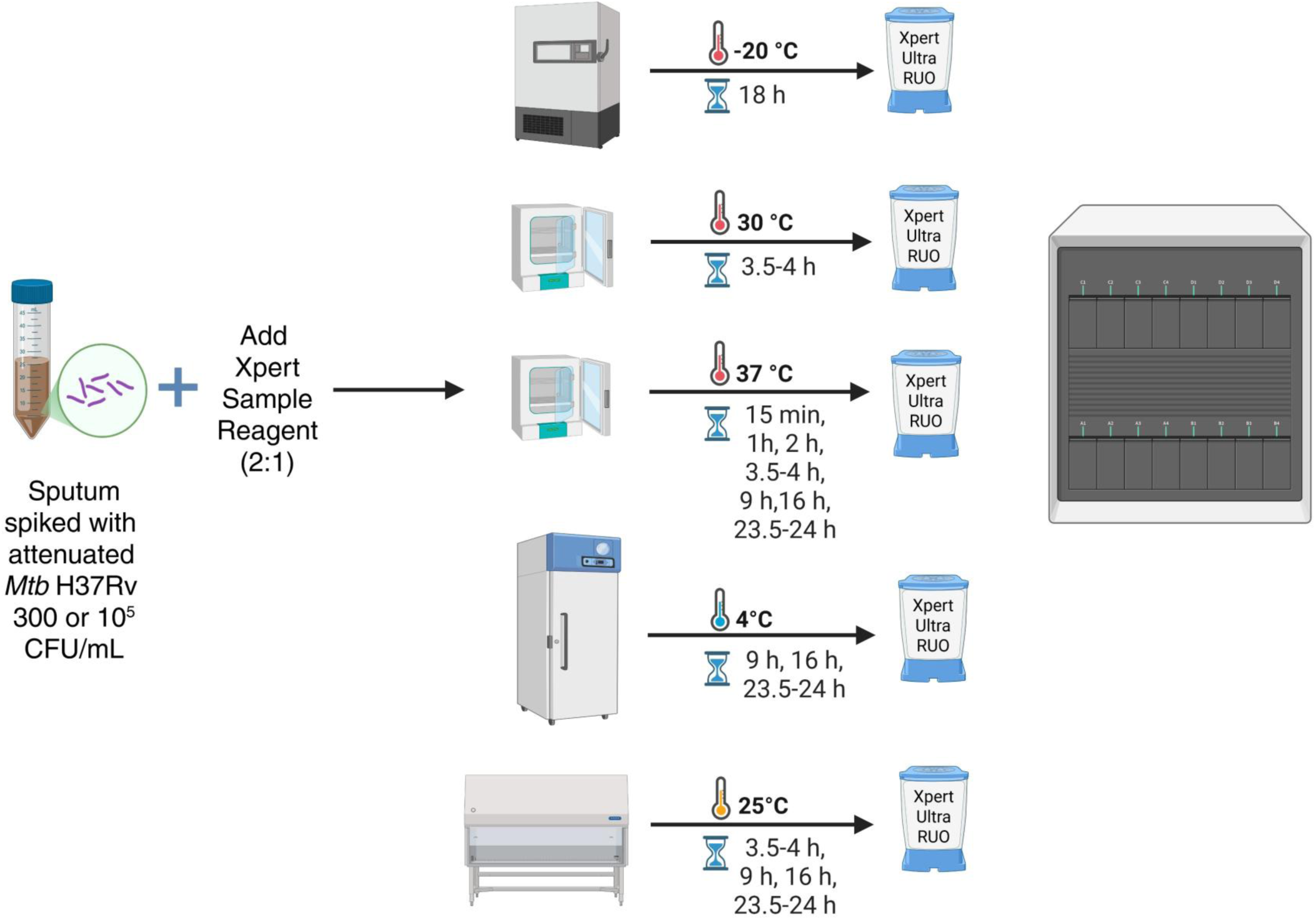
Sample processing workflow showing how the TB-negative sputum spiked with “low” (300 CFU/mL) and “high” (100,000 CFU/mL) bacterial load of *Mycobacterium tuberculosis* H37Rv in Xpert sample reagent (SR) was tested at different temperatures and incubation times.

**Table 1.**
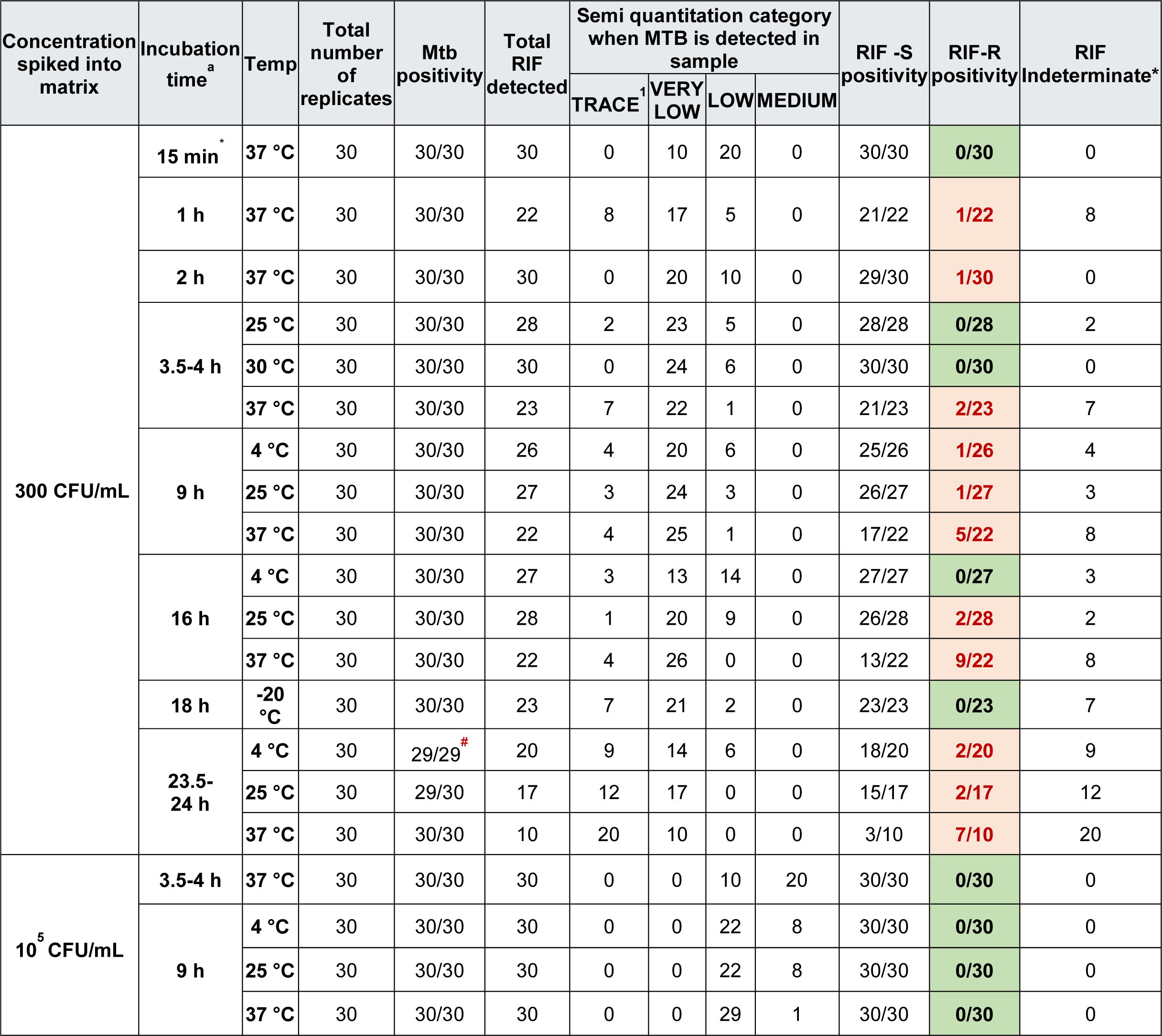
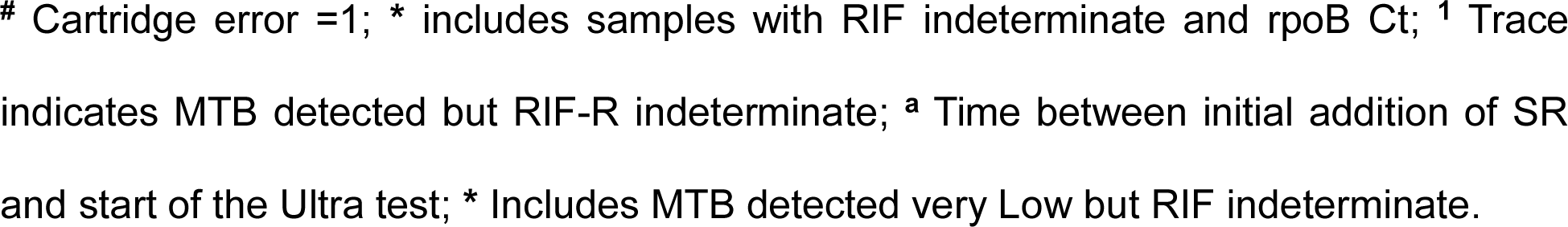
Xpert Ultra performance with “low/very low” (300 CFU/mL) and “medium” (100,000 CFU/mL) of attenuated H37Rv at different temperatures and SR hold time.

| Concentration spiked into matrix | Incubation time<sup>a</sup> | Temp | Total number of replicates | Mtb positivity | Total RIF detected | Semi quantitation category when MTB is detected in sample |  |  |  | RIF -S positivity | RIF-R positivity | RIF Indeterminate* |
| --- | --- | --- | --- | --- | --- | --- | --- | --- | --- | --- | --- | --- |
|  |  |  |  |  |  | TRACE<sup>1</sup> | VERY LOW | LOW | MEDIUM |  |  |  |
| 300 CFU/mL | 15 min<sup>*</sup> | 37 °C | 30 | 30/30 | 30 | 0 | 10 | 20 | 0 | 30/30 | 0/30 | 0 |
|  | 1 h | 37 °C | 30 | 30/30 | 22 | 8 | 17 | 5 | 0 | 21/22 | 1/22 | 8 |
|  | 2 h | 37 °C | 30 | 30/30 | 30 | 0 | 20 | 10 | 0 | 29/30 | 1/30 | 0 |
|  | 3.5-4 h | 25 °C | 30 | 30/30 | 28 | 2 | 23 | 5 | 0 | 28/28 | 0/28 | 2 |
|  |  | 30 °C | 30 | 30/30 | 30 | 0 | 24 | 6 | 0 | 30/30 | 0/30 | 0 |
|  |  | 37 °C | 30 | 30/30 | 23 | 7 | 22 | 1 | 0 | 21/23 | 2/23 | 7 |
|  | 9 h | 4 °C | 30 | 30/30 | 26 | 4 | 20 | 6 | 0 | 25/26 | 1/26 | 4 |
|  |  | 25 °C | 30 | 30/30 | 27 | 3 | 24 | 3 | 0 | 26/27 | 1/27 | 3 |
|  |  | 37 °C | 30 | 30/30 | 22 | 4 | 25 | 1 | 0 | 17/22 | 5/22 | 8 |
|  | 16 h | 4 °C | 30 | 30/30 | 27 | 3 | 13 | 14 | 0 | 27/27 | 0/27 | 3 |
|  |  | 25 °C | 30 | 30/30 | 28 | 1 | 20 | 9 | 0 | 26/28 | 2/28 | 2 |
|  |  | 37 °C | 30 | 30/30 | 22 | 4 | 26 | 0 | 0 | 13/22 | 9/22 | 8 |
|  | 18 h | -20 °C | 30 | 30/30 | 23 | 7 | 21 | 2 | 0 | 23/23 | 0/23 | 7 |
|  | 23.5-24 h | 4 °C | 30 | 29/29<sup>#</sup> | 20 | 9 | 14 | 6 | 0 | 18/20 | 2/20 | 9 |
|  |  | 25 °C | 30 | 29/30 | 17 | 12 | 17 | 0 | 0 | 15/17 | 2/17 | 12 |
|  |  | 37 °C | 30 | 30/30 | 10 | 20 | 10 | 0 | 0 | 3/10 | 7/10 | 20 |
| 10<sup>5</sup> CFU/mL | 3.5-4 h | 37 °C | 30 | 30/30 | 30 | 0 | 0 | 10 | 20 | 30/30 | 0/30 | 0 |
|  | 9 h | 4 °C | 30 | 30/30 | 30 | 0 | 0 | 22 | 8 | 30/30 | 0/30 | 0 |
|  |  | 25 °C | 30 | 30/30 | 30 | 0 | 0 | 22 | 8 | 30/30 | 0/30 | 0 |
|  |  | 37 °C | 30 | 30/30 | 30 | 0 | 0 | 29 | 1 | 30/30 | 0/30 | 0 |

### Retrospective evaluation of historical Xpert Ultra analytical study data

A retrospective analysis was performed using Xpert Ultra test results generated between 2015 and 2025 as part of Technical Feasibility and Design Verification/Validation studies performed at Cepheid and Rutgers. All runs were conducted using strains harboring wild-type *rpoB* gene, namely *M. bovis* BCG, attenuated H37Rv (BSL-2 strain) or external controls with cloned wild-type *rpoB* gene, spiked in either TB-negative sputum or buffer and incubated in SR at times ranging from 15 minutes to 4 hours (10). None of the runs exceeded the recommended SR incubation time of 4 hours. We initially collected “results summary” files from all the Ultra runs performed during this time. We then selected 3835 runs, which generated the assay-defined “MTB DETECTED LOW” or “MTB DETECTED VERY LOW” result outputs, for further analysis. Test runs with “INDETERMINATE”, “ERROR”, “INVALID”, or “MTB DETECTED HIGH” or “MEDIUM” results were excluded.

The objective of this analysis was to specifically identify any instances of false rifampicin resistance (RIF-R) detection in SR-treated, spiked specimens with wild-type strains generating “LOW” and “VERY LOW” semi-quantitation results. Additionally, Ct values and melt-curve data associated with the rifampicin resistance detection probes were reviewed for representative samples across the study period for additional evaluation.

### Library preparation and sequencing analysis with false RIF-R amplicons

Amplified DNA was extracted from the reaction tube of a randomly selected number of false positive Xpert Ultra cartridges with a sterile syringe. The recovered volume was stored in microcentrifuge tubes at - 40^0^C prior to library preparation. A single-step PCR was performed to add adapters to the *rpoB* RRDR region present in the extracted DNA amplicons to generate 200 bp length *rpoB* amplicon libraries for index library preparation with the Illumina Nextera XT index kit v2 (Illumina, San Diego, CA) and libraries sequenced on the Illumina MiSeq (V2, 150 bp paired end chemistry). Sequencing results have been deposited in the NCBI Sequence Read Archive under SRA Project number PRJNA1538067.

Fastp 1.0.1 (11) was used to remove adapter sequences and low-quality reads. Minimap2 (version 2.28) (12) was used to align the trimmed reads to *rpoB* sequence (NCBI Ref Seq AL123456.3:759807-763325). SAMtools (version 1.23) (13) was used to prepare the bam files. The variants in the false positive amplicons were visualized on the Integrative Genomic Browser (IGV) (version 2.16.1) (14). *rpoB* mutations were confirmed with TB-profiler tool (version 6.7.0).

### Statistical analysis

All statistical analyses were performed using GraphPad Prism Version 10.4.2.

## Results

### Frequency of rifampicin susceptible results at “LOW” and “VERY LOW” Xpert Ultra results

We undertook a systematic and comprehensive analysis of the Xpert Ultra runs from the analytical and assay validation experiments performed at the Cepheid and Rutgers laboratories during a period of ten years to see if we ever encountered such false RIF-R results with paucibacillary specimens. For this purpose, we only analyzed the runs which were performed with known *M. bovis* BCG, or attenuated H37Rv (BSL2 version) isolates, or wild-type external control cells (containing a wild-type copy of the *rpoB* gene) spiked in sputum or buffer. These runs were representative of three main categories of analytical R&D studies performed during and after the development of the Xpert Ultra assay; namely relevant technical feasibility and design verification studies and all experiments performed for estimation of limit of detection and bead stability determination (**Fig. 2**). Of the 3835 Xpert Ultra runs tested as part of various analytical studies between 2015 and 2025, a total of 2413 were Ultra “low,” and 1422 were Ultra “very low” results (**Fig 2**). All the runs analyzed, employed SR incubation either for 15 minutes, or no more than 4 hours as specified in the package insert (10). All the 3835 runs correctly generated “RIF Resistance NOT DETECTED” result. Not a single false positive rifampicin resistance (RIF-R) result was detected in any of these runs for any of the samples with any level of semi-quantitative outputs including the “low” and “very low” results, with correct wild-type Tm values generated for all the four rpoB RRDR probes. Additionally, we analyzed the melt profiles of randomly selected 100 test runs to investigate if we could identify any melt curve aberrations which may eventually increase the risk of false RIF-R calls, even though all the results were accurate. We did not observe any alterations or aberrations in the first derivative melt curve profiles which can lead to such false positive RIF-R calls specifically in samples with very low/low bacillary semi-quantitation.

**FIG 2.**
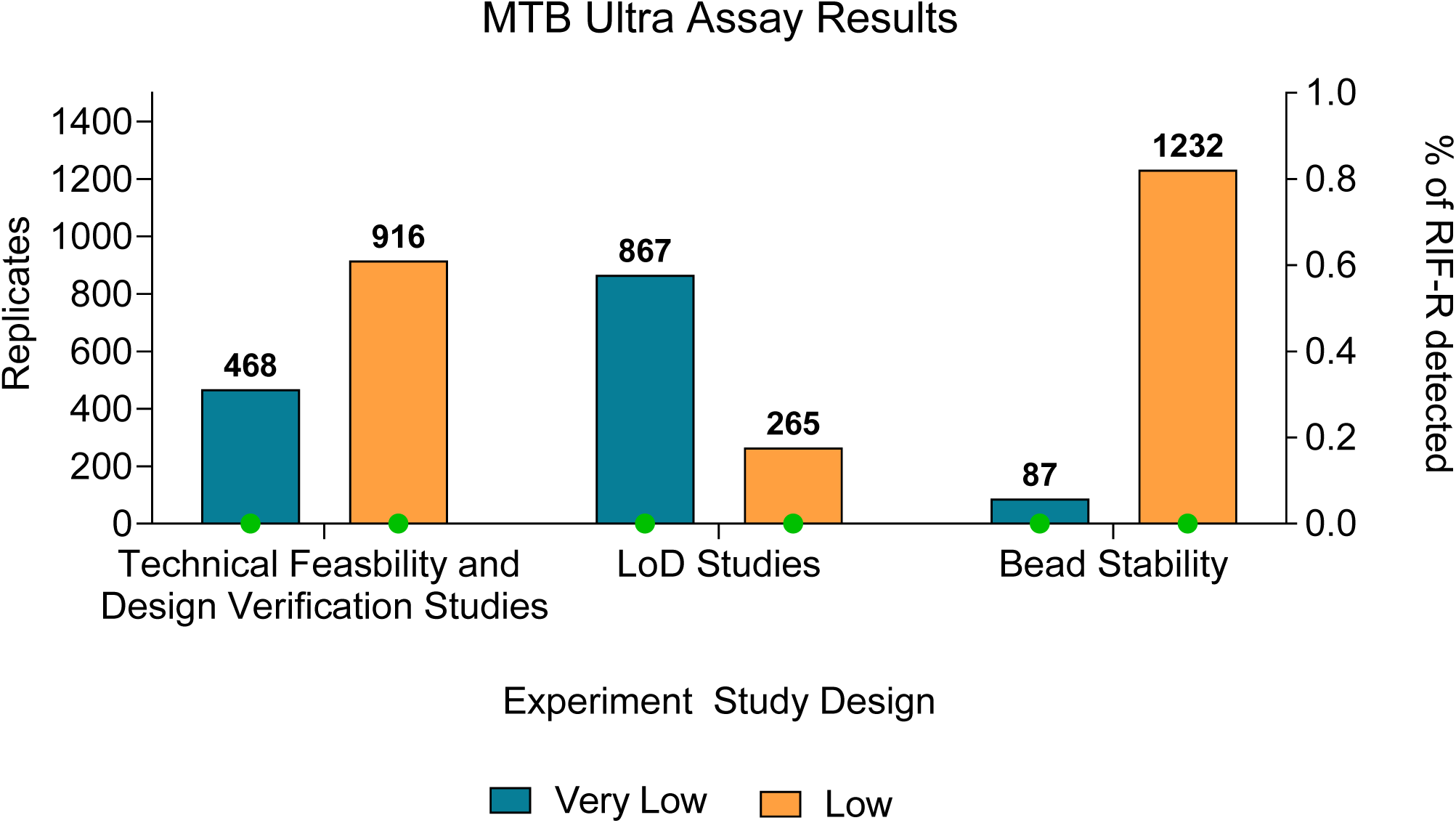
Retrospective analysis of archived analytical data: Graph showing total number of “Low” and “Very low” results obtained from various analytical studies performed over a period of 10 years. A total of 2413 “Low” and 1422 “Very Low” runs using BCG, or attenuated H37Rv (BSL2 version) or wild type external control cells spiked in sputum or buffer are represented in the graph. Not a single false RIF-R call has been obtained thus far in any of these runs. The SR hold times range from 15 minutes to 4 hours at room temperature for all the runs. Green dots indicate 0% of the runs resulting in false RIF-R calls.

### Performance of Xpert Ultra assessed at higher and lower bacterial loads with off-label duration of SR incubation

The manufacturer recommended limit for testing SR treated sputum samples in Xpert Ultra assays is 15 minutes, to 4 hours (10). Here we assessed the effect of prolonged SR incubation on RIF-R detection results at different temperatures (−20°C, 4°C, 25°C, 30°C and 37°C) and different incubation times (ranging from 15 min to 24 h) for contrived sputum samples with higher and lower bacterial loads of *Mtb* H37Rv (**Table 1**). *Mtb* was detected correctly for 598 of a total of 599 runs, excluding 1 “Error”. In the sample set with a higher mycobacterial load (100,000 CFU/mL), Xpert Ultra generated “low” and “medium” semi-quantitation results for 83 (69.17%) and 37 (30.83%) tests respectively (**Table 1**). No false RIF-R results were observed for this sample set at any temperature or SR incubation times tested, irrespective of the duration of the incubation or temperature of incubation even though 69% of the samples generated a “low” semi-quantitation measurement. This indicated that samples with comparatively higher mycobacterial loads (comparable to microscopy smear-positive samples) are resilient to the effects of SR incubation for as long as 9 hours even at 37^0^C, and do not lead to false RIF-R detection (**Table 1**).

Of the 479 valid tests performed with sputum spiked with a lower bacterial load (300 CFU/mL), Ultra estimated the bacterial load as “trace” (84/479, 17.54 %), “very low” (306/479; 63.88%) and “low” (88/479; 18.37%) categories (**Table 1**). We did not observe any false RIF-R results for the test conditions: −20^0^C hold for 18 h, 4^0^C hold for 16 h, 25^0^C and 30^0^C hold for 3.5-4 h, and 37^0^C hold for 15 min. However, we observed multiple cases of false RIF-R results for the runs which generated a valid RIF-result output (other than RIF-Indeterminate) at all the other conditions tested where SR hold time exceeded the recommended duration (**Table 1**, **Fig 3**). Except for the 15 -minute hold time, the highest proportion of false RIF-R results were obtained at 37^0^C, especially with longer hold times (**Table 1**, **Fig 3**); thus, both elevated temperature or/and prolong hold time appear to interact and produce increasing risk of false RIF-R identification.

**FIG 3.**
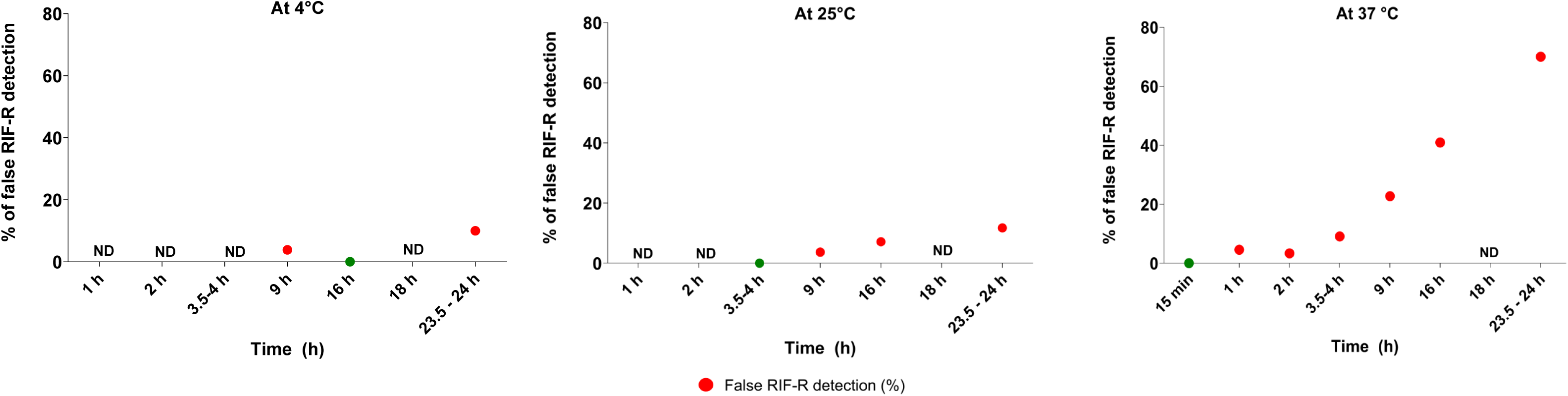
Percentage of false resistance detection in spiked sputum across different incubation times and temperatures with Xpert Sample Reagent (SR) (a) 4 °C, (b) 25°C and (c) 37 °C. X axis represents the time in hours, and y axis represents the % of false RIF-R detection; ND – not done, red dots – false RIF resistance, green dots – absence of false RIF resistance.

### Melt and mutant Tm profile of the test runs generating false RIF-R results

The Xpert MTB/RIF Ultra assay detects a mutation in the RRDR region by a shift in the melting temperature (Tm) of at least one of the 4 sloppy molecular beacons (SMB) from the wild type Tm window (2). We detected 8.5% (33/385) false RIF-R results among all the valid RIF calls from Xpert Ultra runs. Of the 33 false RIF-R results detected in our study: one sample generated mutant Tm values from 3 of the 4 rpoB probes, 6 samples generated mutant Tm values from 2 of the 4 probes, and 26 samples produced mutant Tm value from only 1 of the 4 probes. We observed that Ultra detected RIF-R in 2/33 samples, that generated only 3 Tm values when either the rpoB3 (Tm present for rpoB1 MUT, rpoB2 WT, rpoB4 MUT) or rpoB4 (Tm present for rpoB1 WT, rpoB2 WT, rpoB3 MUT) failed to generate Tm values in both the wild type and mutant windows (**Table S1)**. This is consistent with previous observations on the detection of RRDR mutations with 3 Tm values (15). The mutant melting temperatures (Tm) detected in all 33 false RIF-R samples (**Fig S1**) corresponded to clinically relevant resistance-associated Tm shifts (15).

Ultra resistance calls with a rpoB probe displaying both wild-type, and mutant melting temperature peaks indicate RIF-heteroresistance (2, 8). We observed heteroresistance-like pattern with at least one rpoB probe generating both the mutant Tm and corresponding wild type Tm in 45% (15/33) of the Ultra runs with false RIF-R calls. We observed heteroresistance-like patterns in rpoB1 (10/33), rpoB3 (2/33) and rpoB4 (4/33) with both mutant and wild type Tm values detected in these samples. This specifically indicates on-going SR induced mutagenesis in a wild-type background during prolonged SR incubation (**Fig. 4, Table S1**).

**FIG 4.**
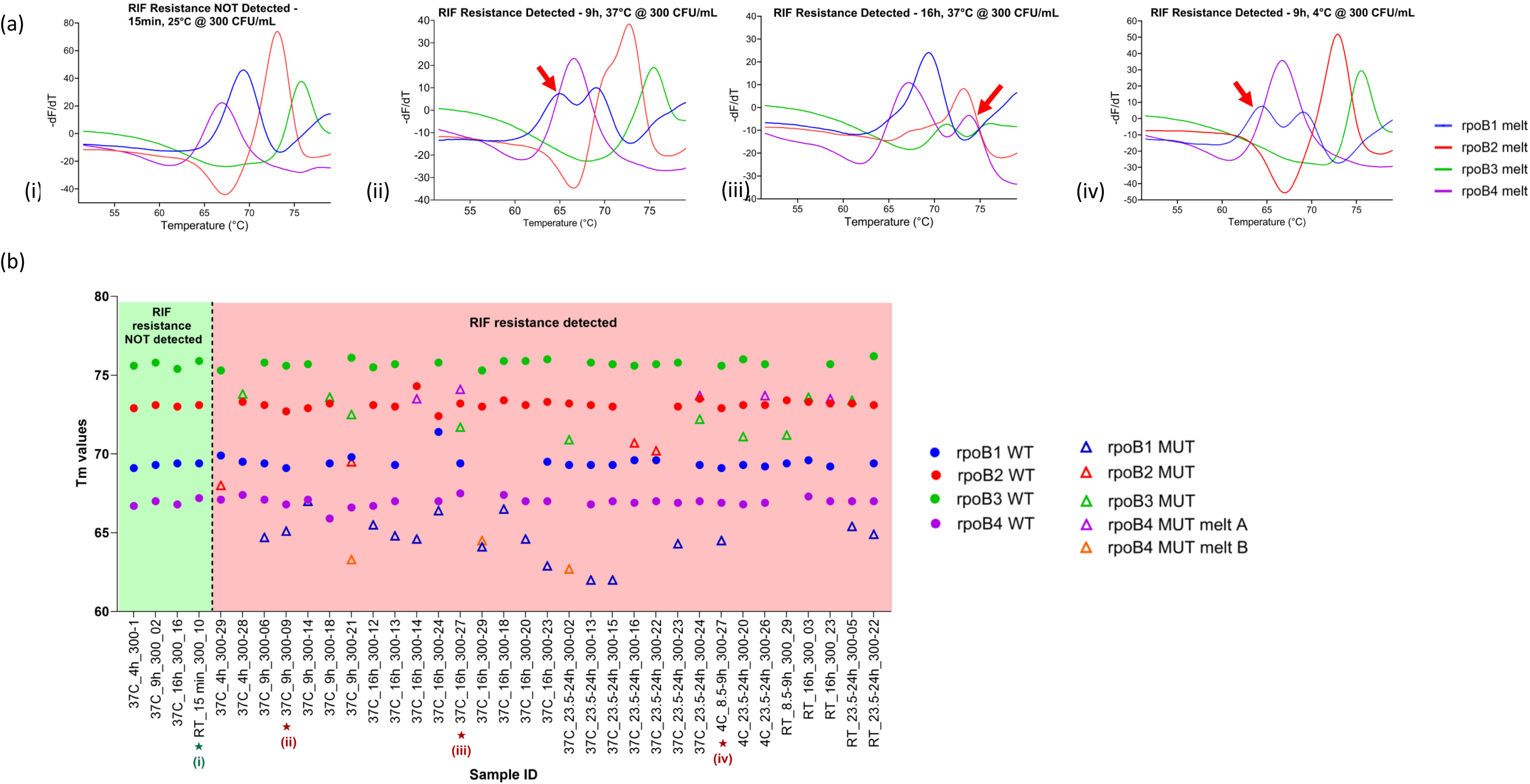
(a) Detection of false heteroresistance in sputum samples spiked with wild type *Mtb* incubated at time points and temperatures deviating from the standard Cepheid Xpert Ultra protocol (b) Tm window pattern of *rpoB* probes in samples designated with “RIF resistance not detected “and “RIF resistance detected” calls.

### Deep sequencing of Ultra amplicons to confirm presence of SR incubation induced

#### *rpoB* RRDR mutations

We sequenced a subset of the rpoB amplicons from low bacterial load samples with false RIF-R calls to confirm if the nucleotide changes occurred due to prolonged SR incubation. TB-profiler (16, 17) identified the presence of mutations due to a nucleotide change from C>T in the *rpoB* amplicon of all the false RIF-R runs tested (**Table S2, Fig S2, Fig S3**). This is consistent with SR-induced deamination. However, these individual mutations are not reported to be phenotypically associated with rifampicin or rifapentine drug resistance (18).

## Discussion

The Ultra assay uses SMB probe chemistry and detects mutations by measuring a characteristic shift in Tm values from at least one of its four melt probes. These SMB Tm values are completely independent of mycobacterial load, or input DNA amount, as demonstrated by several studies which we systematically performed over the years (2, 19-22). In samples containing mixtures of wild-type (WT) and mutant DNA, each sequence type consistently produces its corresponding Tm value, regardless of the relative concentration of either DNA type (varying from 5% to 95% in a mixture) (2). Similarly, in dynamic range studies, all the WT Tm values remain consistent throughout the entire dynamic range from very high to very low bacillary input (2, 19-22). These meticulous studies confirm that the Tm values from all four probes remain reproducible despite differences in the DNA input amount and do not alter or shift with reduced template input. We further confirmed these observations by our retrospective data analysis of more than 10 years of historical Ultra run data, comprising of nearly four thousand tests, which did not identify a single instance where reduced DNA amount or lower bacillary load caused Ultra Tm values to shift from the WT to the mutant Tm window, causing false RIF-R calls. This suggests that it is highly unlikely to get false RIF-R results associated with paucibacillary load samples; at least analytically, especially when SR incubation times are consistent with manufacturer’s instructions.

For any given Ultra SMB probe, a Tm shift from the WT window to a mutant window can occur for three main reasons: (i) a mutation within the probe footprint (whether naturally occurring or artificially induced with a mutating agent) that destabilizes or stabilizes probe binding (23); (ii) DNA “breathing” or hydrogen-bond disruption caused by agents such as excessive detergent, betaine, or glycerol, which lower the probe-target hybrid Tm (23, 24); or (iii) substantially reduced or increased MgCl2, which weakens or strengthens probe-target binding respectively (25, 26). Based on this background and the Ultra assay’s melt-probe chemistry, we analytically investigated what could be the likely cause of the unusually high false RIF-R results reported in paucibacillary specimens in recent clinical studies (5, 27). We reasoned that these false RIF-R results were unlikely to be attributable to low bacillary load alone given our long experience with designing such assays. Given that at least one of the studies (5) mentioned significantly prolonging the SR incubation time beyond the recommended limit, we hypothesized that this could be one of the potential reasons leading to DNA mutagenesis in the sputum samples, and false RIF-R results due to the assay in essence correctly detecting those chemically induced mutations.

Previous studies have shown that incubation in SR for an extended period resulted in false-positive detection of rifampicin resistance in the Xpert® MTB/RIF cartridge due to chemically induced deamination of DNA bases in the *rpoB* RRDR (7). Additionally, it has been shown that temperature speeds up the deamination of nucleic acids in the presence of sodium hydroxide (6). Our systematic analytical study clearly shows that increased false RIF-R occurs at time points beyond 15 minutes at 37^0^C and above 4 hours at lower temperatures. This prolonged SR incubation induced false RIF-R calls can be explained by the SR induced mutagenesis of the Mtb DNA in the samples, which was confirmed by deep sequencing the amplicons which showed mutations consistent with deamination, specifically in the paucibacillary samples with false RIF-R calls. In addition to these false RIF-R calls remaining restricted to the paucibacillary sample similar to the observations in these two studies (5, 27), we also observed an increased frequency of hetero-resistance melt patterns as indicated in one of the studies (27). Investigating the exact reason behind mycobacterial load-dependent SR induced mutagenesis was out of scope of this study. However, the lack of false RIF-R detection in the higher bacterial load samples could be explained if induced SR mutations only develop in a small proportion of the bacterial genomes present in a sample. Under such circumstances, these few mutations might be easily detected when only a small number of total genomes are present, but the mutation signal would be overwhelmed by a vast excess, or wild type genomes present in a sample with a high bacterial load. This occurrence could well explain why false RIF-R calls and increased hetero-resistance pattern were observed only in samples with a lower mycobacterial load when subjected to prolonged SR hold in the clinical study in Rwanda (5).

We have analytically demonstrated that extended incubation of spiked sputum samples in Xpert SR can induce false resistance in Ultra assay, specifically with paucibacillary samples. However, our study is limited by the lack of corresponding confirmation in clinical settings. The Rwanda study (5) estimated the Positive Predictive Value (PPV) for RIF-R detection with a large prospective sample set with 53,096 Ultra results. However, the off-label use of Xpert SR in this study implies that more large-scale clinical studies with paucibacillary samples are needed to determine if Ultra assay would indeed lead to the same issue of increased false RIF-R calls with paucibacillary samples when used according to the manufacturer’s recommendations. Such use of the Ultra test with prolonged SR incubation of sputum samples significantly beyond the specifically recommended time limit in the manufacturer’s instructions for users has not been reviewed by any regulatory body. To the best of our knowledge, apart from these two recent clinical studies(5, 27), there have been no other studies performed in nearly a decade since the Ultra assay was launched, which reported similar issues with the assay. We have analytically reproduced the findings in the Rwanda clinical study showing a clear relationship of prolonged off label SR hold with false RIF-R calls in paucibacillary specimens and believe that such observations will be extremely rare or not observed at all in clinical settings where manufacturer’s instructions on sample processing are adhered to.

## Acknowledgments

The authors thank Cepheid, Inc. (Devasena Gnanashanmugam and Sherri Williams) for donating research-use-only Xpert Ultra cartridges. We thank Elissa Ocke for her advice on the IGV analysis pipeline. The authors acknowledge the Genomics Center Rutgers New Jersey Medical School (https://research.njms.rutgers.edu/genomics/) and the Office of Advanced Research Computing (OARC) at Rutgers, The State University of New Jersey (http://oarc.rutgers.edu) for providing access to the Amarel cluster and associated research computing resources that have contributed to the results reported here.

## Figure Legends

**FIG S1.**
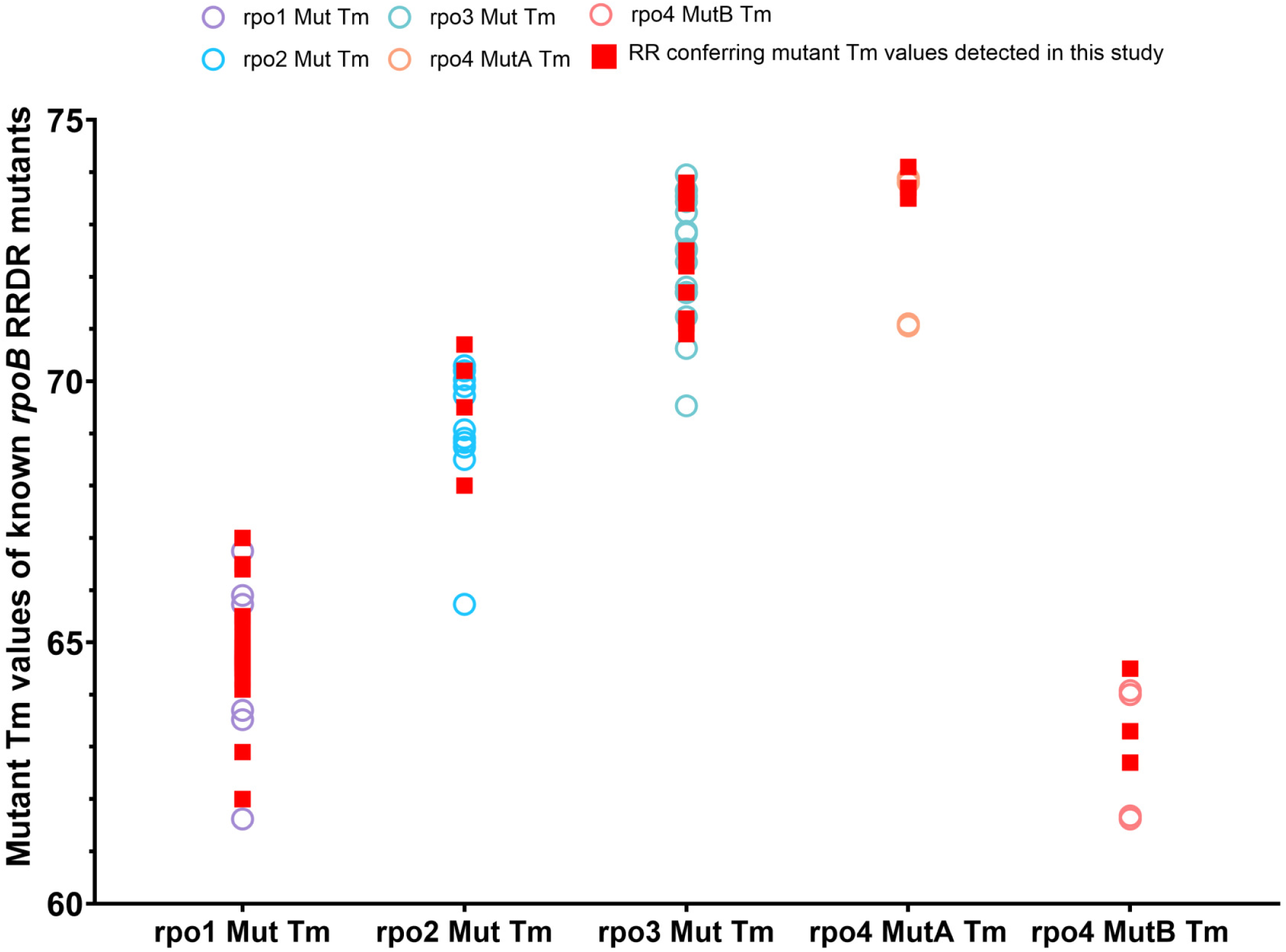
Plot of Ultra Tm values for rpoB probes as observed for samples with false RIF resistance detection calls. Open circles represent Tm of clinical strains (15) and red squares indicate Tm values detected in this study.

**FIG S2.**
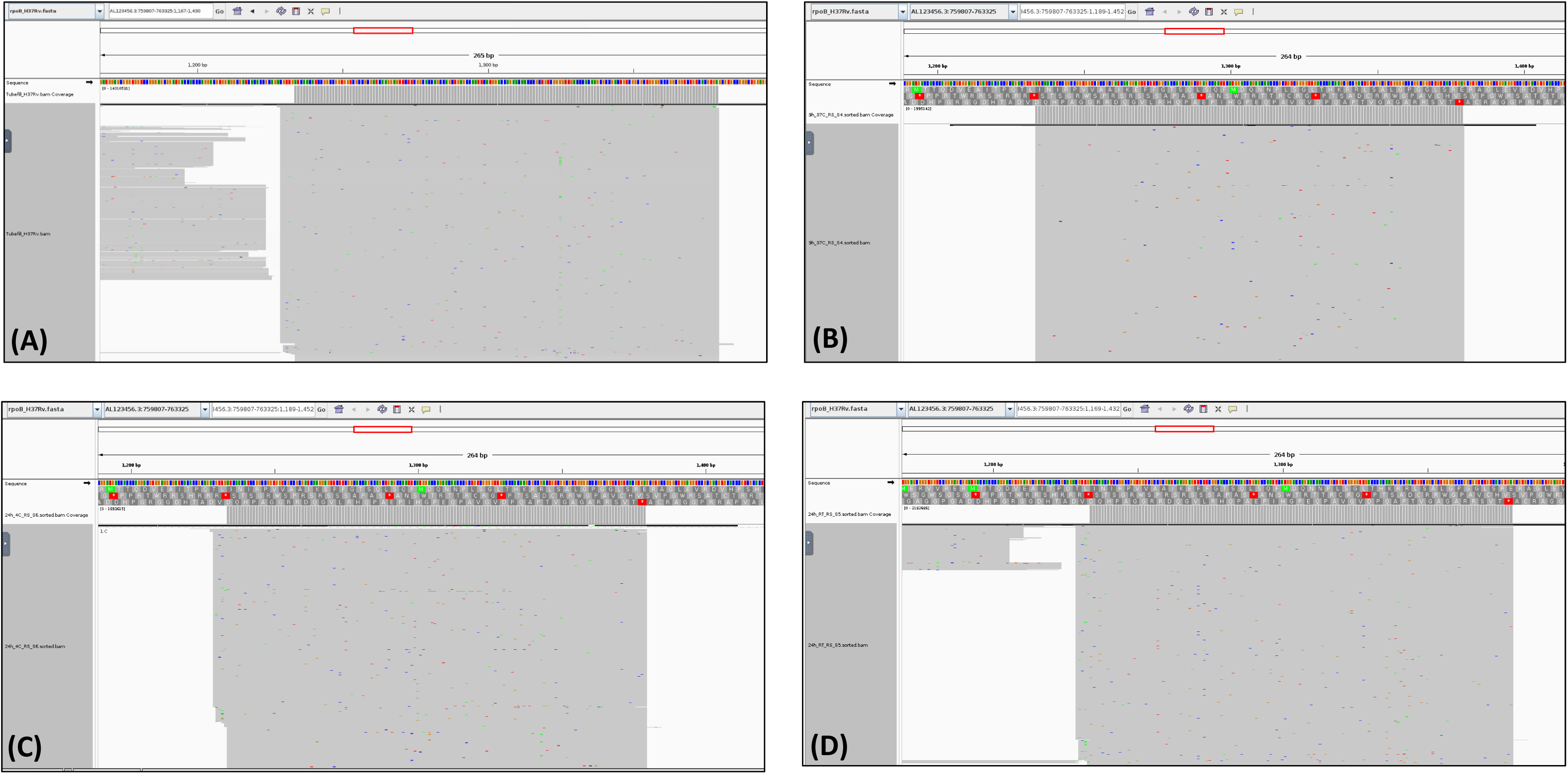
IGV visualization of the RRDR region in samples that generated “RIF-Resistance NOT DETECTED” results with. (A) higher bacterial load – 10^6^ genomic copies and low or very low bacterial load (300 CFU/mL) with sample incubated in SR at (B) 37°C for 9h (C) 4°C for 24h (D) 25°C for 24 h.

**FIG S3.**
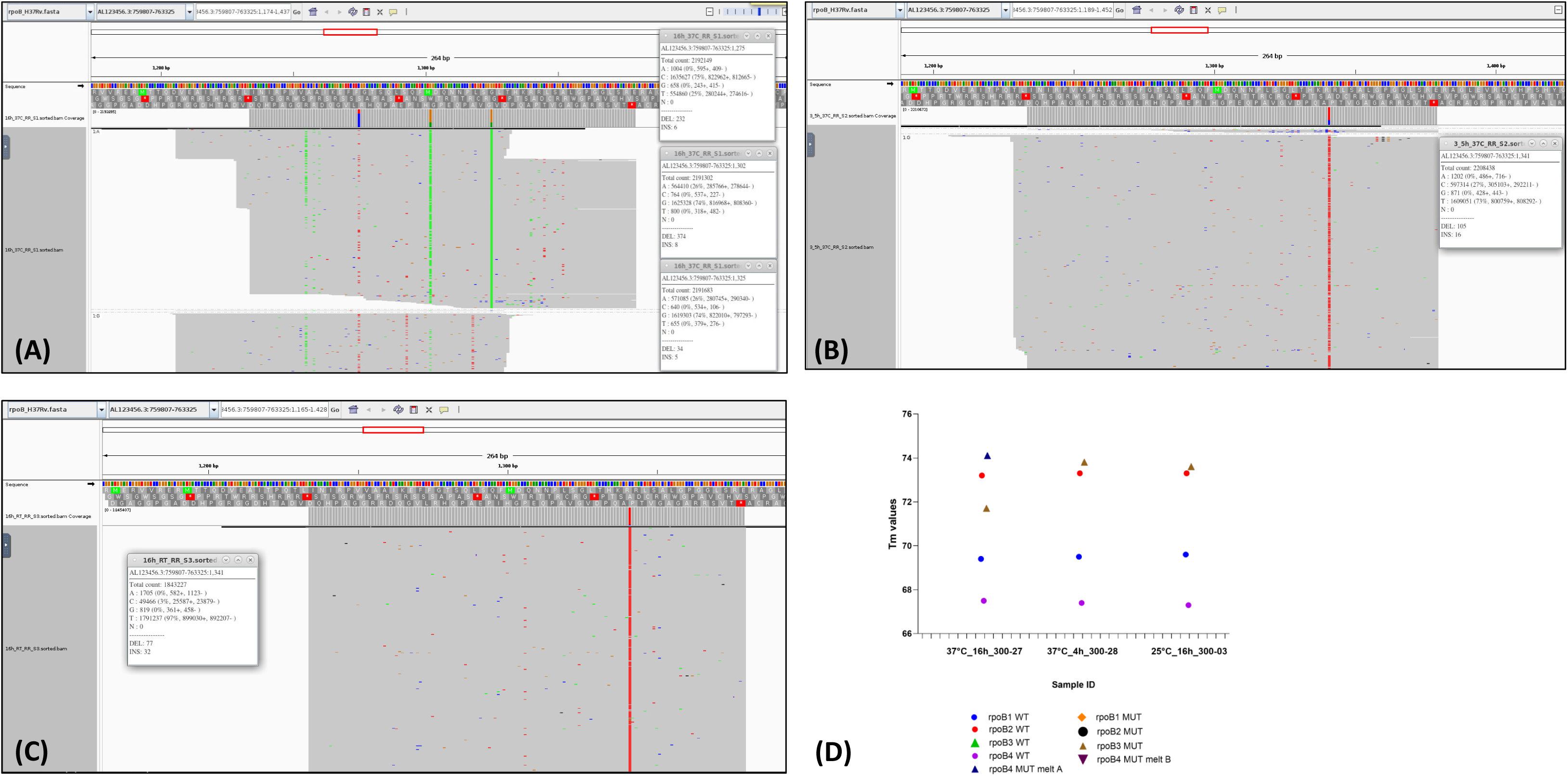
IGV visualization of the RRDR region of low or very low bacterial load (300 CFU/mL) samples that generated “RIF Resistance DETECTED” results when incubated in SR at. (A) 37°C for 16h (B) 37°C for 3.5-4 h (C) 25°C for 16 h and (D) Tm window pattern of *rpoB* probes in samples (A), (B) and (C).

## References

1. World Health Organization. 2025. WHO operational handbook on tuberculosis: module 3: diagnosis.

2. Chakravorty S, Simmons AM, Rowneki M, Parmar H, Cao Y, Ryan J, Banada PP, Deshpande S, Shenai S, Gall A, Glass J, Krieswirth B, Schumacher SG, Nabeta P, Tukvadze N, Rodrigues C, Skrahina A, Tagliani E, Cirillo DM, Davidow A, Denkinger CM, Persing D, Kwiatkowski R, Jones M, Alland D. 2017. The New Xpert MTB/RIF Ultra: Improving Detection of *Mycobacterium tuberculosis* and Resistance to Rifampin in an Assay Suitable for Point-of-Care Testing. mBio 8:10.1128/mbio.00812–17.

3. Haas AL, Ma A, Pham J, Verma P, Malhotra U, Church EC, Narita M, Escuyer V, Shakir SM. 2025. Limitations of the MTB/RIF Assay: An Xpert Review of 4 Clinical Cases. Open Forum Infectious Diseases 12.

4. Lin T-H, Chung H-Y, Jian M, Jr., Chang C-K, Lai Y-W, Perng C-L, Chang F-Y, Chen Y-H, Shang H-S. 2026. Nationwide longitudinal evaluation of a machine learning approach for enhanced interpretation of Xpert MTB/RIF ultra rifampicin-resistance results in low bacterial load tuberculosis specimens. Journal of Infection and Public Health 19:103064.

5. Cuella-Martin I, Hakizayezu F, Ahmed A, Runyambo D, Niyompano H, Keysers J, De Rijk WB, Mulders W, Mitchell EMH, Decroo T, Habimana YM, Migambi P, Muvunyi CM, de Jong BC, Rigouts L, Ngabonziza JCS. 2025. Paucibacillary Tuberculosis Drives the Low Positive Predictive Value of Xpert MTB/RIF Ultra for Rifampicin Resistance Detection in Low-Prevalence Settings. Clinical Infectious Diseases 81:372–378.

6. Wang P, Ren J. 2004. Separation of purine and pyrimidine bases by capillary electrophoresis using β-cyclodextrin as an additive. Journal of Pharmaceutical and Biomedical Analysis 34:277–283.

7. Banada PP, Sivasubramani SK, Blakemore R, Boehme C, Perkins MD, Fennelly K, Alland D. 2010. Containment of bioaerosol infection risk by the Xpert MTB/RIF assay and its applicability to point-of-care settings. J Clin Microbiol 48:3551–7.

8. Rudra P, Parmar H, Daivaa N, Tran C, Narang A, Singh S, Somaiya D, Roybardhan S, Anwesa R, Reiss R, Ward S, Penn-Nicholson A, Vos MD, Armstrong DT, Borkman AL, Ellner J, Dorman SE, Kim S, Yuan L, Alland D, Chakravorty S.A cartridge-based assay for improved detection of multidrug-resistant *Mycobacterium tuberculosis* directly from sputum. Journal of Clinical Microbiology 0:e01100–25.

9. Chilambi GS, Reiss R, Daivaa N, Banada P, Vos MD, Penn-Nicholson A, Alland D. 2025. Optimal processing of tongue swab samples for *Mycobacterium tuberculosis* detection by the Xpert MTB/RIF Ultra assay. Microbiology Spectrum 13:e02403–24.

10. Cepheid. Sunnyvale C, USA. 2019. Xpert® MTB/RIF Ultra (Package insert).

11. Chen S. 2025. fastp 1.0: An ultra-fast all-round tool for FASTQ data quality control and preprocessing. iMeta 4:e70078.

12. Li H. 2018. Minimap2: pairwise alignment for nucleotide sequences. Bioinformatics 34:3094–3100.

13. Li H, Handsaker B, Wysoker A, Fennell T, Ruan J, Homer N, Marth G, Abecasis G, Durbin R, Subgroup GPDP. 2009. The Sequence Alignment/Map format and SAMtools. Bioinformatics 25:2078–2079.

14. Freese NH, Norris DC, Loraine AE. 2016. Integrated genome browser: visual analytics platform for genomics. Bioinformatics 32:2089–2095.

15. Cao Y, Parmar H, Simmons AM, Kale D, Tong K, Lieu D, Persing D, Kwiatkowski R, Alland D, Chakravorty S. 2019. Automatic Identification of Individual *rpoB* Gene Mutations Responsible for Rifampin Resistance in Mycobacterium tuberculosis by Use of Melting Temperature Signatures Generated by the Xpert MTB/RIF Ultra Assay. Journal of Clinical Microbiology 58:10.1128/jcm.00907–19.

16. Coll F, McNerney R, Preston MD, Guerra-Assunção JA, Warry A, Hill-Cawthorne G, Mallard K, Nair M, Miranda A, Alves A, Perdigão J, Viveiros M, Portugal I, Hasan Z, Hasan R, Glynn JR, Martin N, Pain A, Clark TG. 2015. Rapid determination of anti-tuberculosis drug resistance from whole-genome sequences. Genome Medicine 7:51.

17. Phelan JE, O’Sullivan DM, Machado D, Ramos J, Oppong YEA, Campino S, O’Grady J, McNerney R, Hibberd ML, Viveiros M, Huggett JF, Clark TG. 2019. Integrating informatics tools and portable sequencing technology for rapid detection of resistance to anti-tuberculous drugs. Genome Medicine 11:41.

18. World Health Organization. 2023. Catalogue of mutations in Mycobacterium tuberculosis complex and their association with drug resistance, second edition..

19. Chakravorty S, Roh SS, Glass J, Smith LE, Simmons AM, Lund K, Lokhov S, Liu X, Xu P, Zhang G, Via LE, Shen Q, Ruan X, Yuan X, Zhu HZ, Viazovkina E, Shenai S, Rowneki M, Lee JS, Barry CE, 3rd, Gao Q, Persing D, Kwiatkawoski R, Jones M, Gall A, Alland D. 2017. Detection of Isoniazid-, Fluoroquinolone-, Amikacin-, and Kanamycin-Resistant Tuberculosis in an Automated, Multiplexed 10-Color Assay Suitable for Point-of-Care Use. J Clin Microbiol 55:183–198.

20. Roh SS, Smith LE, Lee JS, Via LE, Barry CE, 3rd, Alland D, Chakravorty S. 2015. Comparative Evaluation of Sloppy Molecular Beacon and Dual-Labeled Probe Melting Temperature Assays to Identify Mutations in Mycobacterium tuberculosis Resulting in Rifampin, Fluoroquinolone and Aminoglycoside Resistance. PLoS One 10:e0126257.

21. Chakravorty S, Kothari H, Aladegbami B, Cho EJ, Lee JS, Roh SS, Kim H, Kwak H, Lee EG, Hwang SH, Banada PP, Safi H, Via LE, Cho SN, Barry CE, 3rd, Alland D. 2012. Rapid, high-throughput detection of rifampin resistance and heteroresistance in Mycobacterium tuberculosis by use of sloppy molecular beacon melting temperature coding. J Clin Microbiol 50:2194–202.

22. Chakravorty S, Aladegbami B, Thoms K, Lee JS, Lee EG, Rajan V, Cho EJ, Kim H, Kwak H, Kurepina N, Cho SN, Kreiswirth B, Via LE, Barry CE, 3rd, Alland D. 2011. Rapid detection of fluoroquinolone-resistant and heteroresistant Mycobacterium tuberculosis by use of sloppy molecular beacons and dual melting-temperature codes in a real-time PCR assay. J Clin Microbiol 49:932–40.

23. Miranda P, Weber G. 2021. Thermodynamic evaluation of the impact of DNA mismatches in PCR-type SARS-CoV-2 primers and probes. Molecular and Cellular Probes 56:101707.

24. Song C, Castellanos-Rizaldos E, Bejar R, Ebert BL, Makrigiorgos GM. 2015. DMSO Increases Mutation Scanning Detection Sensitivity of High-Resolution Melting in Clinical Samples. Clin Chem 61:1354–62.

25. Rejali NA, Ye FD, Zuiter AM, Keller CC, Wittwer CT. 2021. Nearest-neighbour transition-state analysis for nucleic acid kinetics. Nucleic Acids Res 49:4574–4585.

26. Lorenz TC. 2012. Polymerase chain reaction: basic protocol plus troubleshooting and optimization strategies. J Vis Exp doi:10.3791/3998:e3998.

27. Ghebrekristos Y, Ahmed A, Beylis N, Singh S, Opperman C, Naufal F, Folkerts M, Engelthaler D, Auma E, Venter R, Booley G, Metcalfe J, Warren R, Theron G. 2025. Xpert MTB/RIF Ultra-resistant and MTBDR *plus* susceptible rifampicin results in people with tuberculosis: utility of FluoroType MTBDR and deep sequencing. Antimicrobial Agents and Chemotherapy 69:e01671–24.

